# The roles of dispersal limitation and pre-adaptation in shaping *Paraburkholderia* endosymbiont frequencies in social amoeba communities

**DOI:** 10.1101/2025.05.20.654931

**Authors:** James G. DuBose, Terry Uhm, Jordan Bowen, Patricia Fiedorek, Mackenzie Hoogshagen, Tamara S. Haselkorn, Susanne DiSalvo

**Author notes:** Denotes co-senior authorship (Tamara S. Haselkorn, Susanne DiSalvo).

## Abstract

Endosymbiotic interactions have long played fundamental roles in shaping the evolution and diversification of eukaryotes. However, we still have a limited understanding of how ecological processes govern the distribution of endosymbionts that are still segregating in host populations. To contribute to this understanding, here we use the interactions between *Paraburkholderia* endosymbionts and their Dictyostelid social amoeba hosts as a model system to investigate the role of dispersal, a fundamental ecological process, in shaping the distribution and evolution of endosymbiotic interactions. We first found that patterns of endosymbiont diversification were highly biogeographic, suggesting a significant degree of dispersal limitation. We then experimentally mediated the dispersal of several endosymbiont species into environments with multiple host species and found that each symbiont was able to sustain a high prevalence in each host population. The benefit/detriment of these mediated interactions did not change with increasing phylogenetic distance from what is suspected to be the focal amoeba host species in nature. Taken together, our findings suggest *Paraburkholderia* endosymbionts are generally pre-adapted to occupy a variety of Dictyostelid host environments, and their distribution among host populations is subject to a high degree of dispersal limitation. Overall, our findings have significant implications for our understanding of how ecological processes facilitate and limit the evolution of endosymbiotic interactions.

**Importance:** Endosymbiotic interactions are ubiquitous in complex eukaryotes, as organelles such as mitochondria and chloroplasts represent the remnants of what were once free-living prokaryotes. However, how ecological processes facilitate the transition from free-living to host-associated is less understood. Selection is the most commonly invoked process to explain this transition: symbionts that are better at infecting hosts and potentially confer some benefit rise in frequency because they are selected for (and otherwise selected against). However, this only describes one fundamental process that can shape the ecology of symbiotic interactions. Here we present evidence that the importance of dispersal (and its limitations) likely exceeds that of selection in shaping the distribution and frequency of *Paraburkholderia* endosymbionts in their Dictyostelid social amoeba host communities. These findings highlight the need to consider the interaction between ecological processes, rather than individual processes, when understanding the ecology and evolution of endosymbiotic interactions.

## Introduction

It is difficult to understand the evolution of complex eukaryotes without considering the endosymbiotic interactions they have formed and maintained throughout natural history. The most notable examples of this point are mitochondria and chloroplast, which have fundamentally shaped the evolution of eukaryotes (Lynn Margulis 1970). More generally, other endosymbioses have independently arisen that have significantly shaped the evolution and diversification of many lineages (Jeon and Jeon 1976; Baumann et al. 1997; Werren 1997; Saikkonen et al. 1998). Notable progress has been made in understanding the molecular and genetic bases underlying a symbiont’s evolutionary transition to an endosymbiotic lifestyle, as well as the macroevolutionary significance to their host’s lineage (McCutcheon and Moran 2012). However, how ecological processes govern the distribution of endosymbionts that are still segregating in host populations has received relatively less empirical attention.

Ecological theory describes the process that shape the distributions and frequencies of organisms as analogous to those that shape the distributions and frequencies of alleles: selection, drift, dispersal (migration), and diversification (mutation) (Vellend 2010). From both an ecological and evolutionary perspective, the primary process ascribed to explain the distribution of symbionts among host taxa is selection: selective pressures exerted by the host facilitates or eliminates symbiont residence, which increases host fitness (Sachs et al. 2011). However, ecological theory recognizes the interplay between different processes and aims to understand the relative importance of each processes in shaping the distributions of organisms. Only recently has investigation into the distribution of host-symbiont interactions begun to shift from a selection-centric framework to one that focuses on integrating more general ecological processes, such as ecological drift and dispersal (Chen et al. 2024).

Despite the prevailing emphasis on host selection, central theory on the evolution of endosymbiotic interactions makes implicit reference to the interplay between dispersal and selection in shaping endosymbiont distributions. The evolutionary training grounds hypothesis suggests that frequent predation by smaller and local eukaryotes (such as amoeba) selects for prokaryotes that can survive intracellular digestion, which then increases their ability to evade immune defenses and infect other hosts (Greub and Raoult 2004; Molmeret et al. 2005). In the context of more general theory: frequent predation by local eukaryotes facilitates pre-adaptation to survive and proliferate in intra-host environments, which confers greater success when a prokaryote disperses to (infects) a new host. Under this interpretation, limitations on dispersal to new host populations, rather than long-term co-evolutionary dynamics, can be a primary driver endosymbiont distribution given significant pre-adaptation. This point has been discussed but has received relatively less attention as a framework for understanding the distribution of symbionts among hosts than long-term co-evolutionary frameworks (Janzen 1980).

Here, we used the interactions between Dictyostelid social amoebae hosts and their *Paraburkholderia* endosymbionts as a model system to investigate the roles of pre-adaptation and dispersal limitation in shaping the distribution of endosymbionts that are segregating in host populations. We first conducted a field study that showed patterns of endosymbiont diversification were highly biogeographic, suggesting an important role of dispersal limitation in shaping their distributions. We then experimentally mediated endosymbiont dispersal to host populations along a continuum of phylogenetic divergence from their focal host and found that each *Paraburkholderia* endosymbiont was able to establish and sustain a high prevalence in each host population. Underlying these interactions, we also found that the fitness consequences (for both host and endosymbiont) did not vary with phylogenetic distance from the focal host, suggesting *Paraburkholderia* endosymbionts are pre-adapted to other hosts that they do not frequently interact with.

## Methods

### Study system

Dictyostelid social amoebae are soil-dwelling protists that prey on bacteria. When local food bacteria are depleted, thousands of individual amoebae cells aggregate to form a multicellular fruiting body that facilitates spore dispersal (Bozzaro 2019). Bacteria that have either stuck to the outside of amoeba cells or survived intracellular digestion are able to persist within these fruiting bodies, thus facilitating their co-dispersal as well (Brock et al. 2011; DiSalvo et al. 2015; Haselkorn et al. 2019). A diversity of bacterial symbionts have been found to associate with Dictyostelid hosts (Sallinger et al. 2021). However, frequency of association, fitness consequences, intracellular dynamics, and genomic evidence of host adaptation has been predominately characterized in three *Paraburkholderia* symbionts whose native amoeba host is thought to be *Dictyostelium discoideum*: *P. agricolaris, P. bonniea*, and *P. hayleyella* (Haselkorn et al. 2019; Brock et al. 2020; DuBose et al. 2022). Therefore, here we focused on studying the ecological processes that shape the distribution and frequency of these three symbionts in Dictyostelid amoeba host communities. For experimental studies, we used *D. discoideum, D. citrinum, D. giganteum, D. purpureum, Polysphondylium violaceum*, and *Cavenderia aureostipes*, because they occur sympatrically and represent a breadth of phylogenetic divergence from *D. discoideum*. For example, *D. citrinum* is a closely related sister species of the same genus, *D. giganteum* and *D. purpureum* are more distantly related members of the same genus, *P. violaceum* is of a different genus in the same family, and *C. aureostipes* is of a different order but in the same clade (Sheikh et al. 2018).

### Characterizing Burkholderiales symbiont prevalence and diversity in natural Dictyostelid social amoeba communities

To study the distribution and frequency of *Paraburkholderia* symbionts in natural Dictyostelid populations, we isolated amoeba from soil collected across the Southeastern United States (Figure 1). We collected each soil sample from just below leaf litter on forest floors or from decaying material in rotting logs and stumps. To obtain amoeba isolates, we plated each soil sample on hay agar (Appendix section 1.1.1) no later than 96 hours after collection. After sufficient time for fruiting body formation, we identified amoeba species based on fruiting body morphology (Raper 2016). Since proliferation of single amoeba cells can give rise to patches of several fruiting bodies, we collected clonal amoeba isolates from 2-5 sori from each patch. We suspended isolate sori in 250 μL of KK2 spore buffer (Appendix section 1.1.3) and extracted total DNA using a Chelex/proteinase K protocol described in (Haselkorn et al. 2019). To confirm that isolates were Dictyostelid fruiting bodies and that DNA extraction was successful, we first conducted a PCR screen on all samples using Dictyostelid-specific primers (D307F and D862R) that target a portion of the 18S rRNA gene (Appendix section 1.2.1) (Baldauf et al. 2018). We excluded samples that did not show amplification, as this indicates the DNA extraction failed, or the sampled fruiting body was not a Dictyostelid. To further confirm the identity of Dictyostelid isolates, we sent a subset of positive PCR products for sequencing Eurofins Genomics (Louisville, KY, USA). We then trimmed sequences using Geneious (v8) (Kearse et al. 2012) and classified isolates by using the NCBI BLAST web application (https://blast.ncbi.nlm.nih.gov/Blast.cgi) to search for matches in the standard nucleotide database.

**Figure 1.**
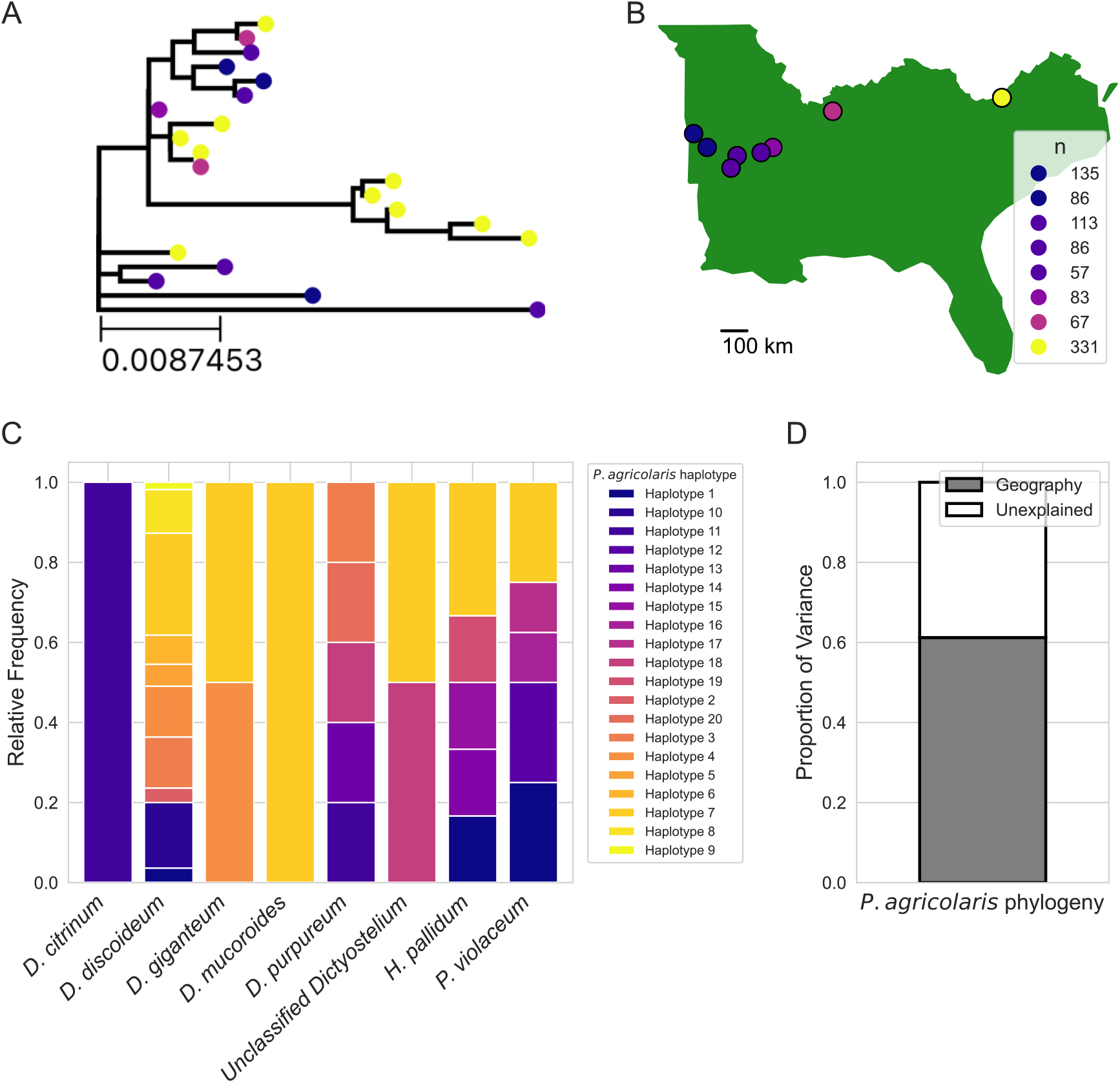
Patterns of *P. agricolaris* divergence are explained by geographic variation, but not patterns of host association. A) The phylogenetic tree of *P. agricolaris* haplotypes found in our surveys of natural populations. Colored circles at the tips indicate the geographic location each haplotype was found in. Note that only a single haplotype was found in multiple (2) locations. B) A map showing the geographic distribution of sampling locations. The legend indicates the number of amoebae isolates collected from each location. C) Stacked bar plots showing the relative frequency of each *P. agricolaris* haplotypes in each social amoeba host. D) A bar graph showing the proportion of variation in patterns of *P. agricolaris* divergence that could be explained by their geographic distribution. Patterns of host association (the frequency of *P. agricolaris* in each host populations) were included in the model, but did not significantly explain any variation.

To identify the presence of Burkholderiales symbionts in each amoeba isolate, we first conducted a PCR screen using Burkholderiales-specific primers (Burk3F and Burk3R) that amplify a portion of the 16S rRNA gene (Appendix section 1.2.2) (Salles et al. 2002). For all positive samples, we then amplified a 780 base-pair region of the leader peptidase A (LepA) gene, which allows for greater species-level identification and haplotype resolution of Burkholderiales symbionts (Haselkorn et al. 2019). We sequenced and identified Burkholderiales symbionts as previously described for the Dictyostelids. We also included the prevalence and LepA sequence data from (DuBose et al. 2022) in our dataset, which allowed us to capture greater geographic and phylogenetic variation in our analyses. After sequence processing, we used MEGA (v7) to align sequences (using the MUSCLE algorithm) and construct a maximum-likelihood phylogenetic tree, where our data was best described by a GTR+G+I substitution model. We used 1000 bootstrap replicates to estimate branch support (Kumar et al. 2016). The only previously described *Paraburkholderia* symbiont we found in nature was *P. agricolaris*. Therefore, we constructed a separate phylogenetic tree using only *P. agricolaris* LepA sequences, where our data was best described by a Tamura 3-parameter+G+I substitution model.

### Experimentally mediating symbiont dispersal to laboratory host populations

To experimentally facilitate the interaction between Dictyostelid hosts and the focal *Paraburkholderia* symbionts, we first plated 10^5^ spores from each tested strain with on 300 μL of a mixture containing 95% *Klebsiella pneumoniae* and 5% symbiont (either *P. agricolaris, P. hayleyella*, or *P. bonniea*, each of which were isolated from *D. discoideum*). We used RFP-labeled symbiont strains because their presence in a fruiting body can be directly visualized via pink discoloration in their colonies upon culturing, and we wanted to perform subsequent microscopy to confirm intracellular infection status (described below) (DiSalvo et al. 2015). We cultured four replica populations for each amoeba and symbiont combination. To examine whether potential symbiont infections persisted beyond initial introduction, we passaged 10 representative fruiting bodies from each host population onto a new plate containing only the host food bacteria *K. pneumoniae*. For passaging, we suspended fruiting bodies in 500 μL of KK2 spore buffer and plated 100 μL of these suspended spores with 200 μL of 1.5 OD_600_ *K. pneumoniae*. We performed two rounds of passaging (for a total of three host generations). Each generation, we measure *Paraburkholderia* prevalence by culturing bacteria from 10 individual host sori, where pink colony discoloration indicated *Paraburkholderia* presence.

### Quantifying intracellular symbiont infection and effects on fitness

To assess fitness cost and and intracellular prevalence of each symbiont-amoeba pairing, we first plated 10^5^ amoeba spores on SM/5 agar plates with 200 μL of *K. pneumoniae* for uninfected references or mixed bacterial suspension consisting of 95% *K. pneumoniae* and 5% RFP-labeled symbiont (adjusted to 1.5 OD600nm). We incubated plates at 24°C for 1 week to allow for fruiting body formation. We then collected and suspended fruiting bodies from the entire plate in in KK2 spore buffer, and quantified total spore productivity via hemocytometer counts. We quantified intracellular infection prevalence by sampling suspended spore samples through a BD-C6 Flow Cytometer, gating spores, and quantifying percent of RFP-positive spores via PE-A intensity histograms, where uninfected samples were used to establish fluorescent and non-fluorescent spore boundaries. To visualize infections via confocal microscopy, we suspended spores from developed fruiting bodies in KK2 buffer supplemented with 1% calcofluor, placed them onto glass-bottom culture dishes, and overlaid 2% agarose. We used an Olympus Fluoview FV1000 confocal microscope equipped with a Plan Apo Oil 60x/1.4 NA objective to acquire images, where we used the DAPI channel to visualize Calcofluor-stained structures (pseudo-colored gray) and the Cy3 channel to visualize RFP (pseudo-colored red). We acquired Z-stacks at 0.5 μm intervals at a resolution of 1024 × 1024 pixels and used Fiji (NIH; imagej.net/Fiji) to process images, including generation of single slices and z-projections.

### Statistics

To quantify the importance of host association frequency and geography in explaining patterns of *P. agricolaris* diversification, we performed a distance-based redundancy analysis using the *capscale* function from the *vegan* R library (Oksanen et al. 2022). First, we used the *cophenetic*.*phylo* function from the *ape* R library to transform the LepA *P. agricolaris* phylogeny into a pairwise phylogenetic distance matrix (Paradis and Schliep 2019) and the *distm* function from the *geosphere* R library to calculate the geographic (Haversine) distance matrix between sample location (Robert J. Hijmans 2023). We then calculated the proportion of variance explained by geographic dissimilarity and host association frequency using the *varpart* function form the *vegan* R library (Oksanen et al. 2022). In addition to a distance-based redundancy analysis, we also performed a Mantel test (with 999 permutations) to quantify the correlation between phylogenetic distance and geographic distance between *P. agricolaris* haplotypes (Oksanen et al. 2022).

To investigate the relationship between phylogenetic distance from *D. discoideum* and fitness effects associated with symbiont infection, we first used the previously described Dictyostelid 18S sequences to create a phylogenetic tree of the social amoebae species used in our study, where our data was best described by a GTR+G+I substitution model. We then quantified the cophenetic distance (which we refer to as phylogenetic distance for simplicity) from each species to *D. discoideum* using the *cophenetic*.*phylo* function from the *ape* R library (Paradis and Schliep 2019). To model the relationship between phylogenetic distance from *D. discoideum* and symbiont intracellular prevalence, we fit a generalized linear model using the *glm* R function with a quasibinomial distribution to account for observed overdispersion (R Core Team 2022). To model the relationship between phylogenetic distance from *D. discoideum* and amoeba fitness effects, we fit a linear model using the *lm* R function (R Core Team 2022).

## Results

### Biogeographic patterns of Paraburkholderia symbionts are strong in Dictyostelid social amoebae populations

To begin understanding the relative importance of dispersal limitation and specific associations in shaping the distribution of *Paraburkholderia* symbionts among social amoeba hosts, we modeled patterns of *Paraburkholderia* phylogenetic divergence (Figure 1A) as a function of their geographic distribution (Figure 1B) and frequencies in different host populations (Figure 1C). We found that the geographic distribution of *P. agricolaris* (the only previously described symbiont found) explained 61.216% of their patterns of phylogenetic divergence (F = 3.1589, p = 0.038) (Figure 1D). However, specific patterns of host association did not significantly explain any of the patterns in *P. agricolaris* phylogenetic divergence (F = 0.9910, p = 0.507), and we did not find any covariance between geographic distributions and specific patterns of host association (Figure 1D). Furthermore, we found a weak but significantly positive correlation between geographic distance and phylogenetic distance between *P. agricolaris* haplotypes (r_M_ = 0.1826, p = 0.012), suggesting a combination of isolation by distance as well as more localized isolation.

### Paraburkholderia symbionts maintain stable associations with a diversity of social amoebae hosts when dispersal is mediated

Our survey of natural social amoebae populations suggest an important role of dispersal limitation in shaping the distribution of *Paraburkholderia* endosymbionts among hosts. Therefore, the following prediction would be that when dispersal is experimentally mediated, *Paraburkholderia* symbionts should be able to establish and persist in various social amoebae populations. To test this prediction, we experimentally introduced the three previously described *Paraburkholderia* symbionts into six different social amoebae populations and measured their prevalence over three generations. Symbiont prevalence remained at or near 100% across generations for most host-symbiont pairings (Figure 2A). The only exception was *P. agricolaris* in *C. aureostipes* hosts, which still maintained 60%-80% prevalence (Figure 2A). This showed that each symbiont was able to stably infect each host population at a high prevalence. Because *Paraburkholderia* symbionts can intracellularly and extracellularly infect amoeba hosts, we used RFP-labeled symbionts and confocal microscopy to confirm that symbionts were intracellularly infecting hosts. This showed that each symbiont was able to infect the spores of each host (Figure 2B).

**Figure 2.**
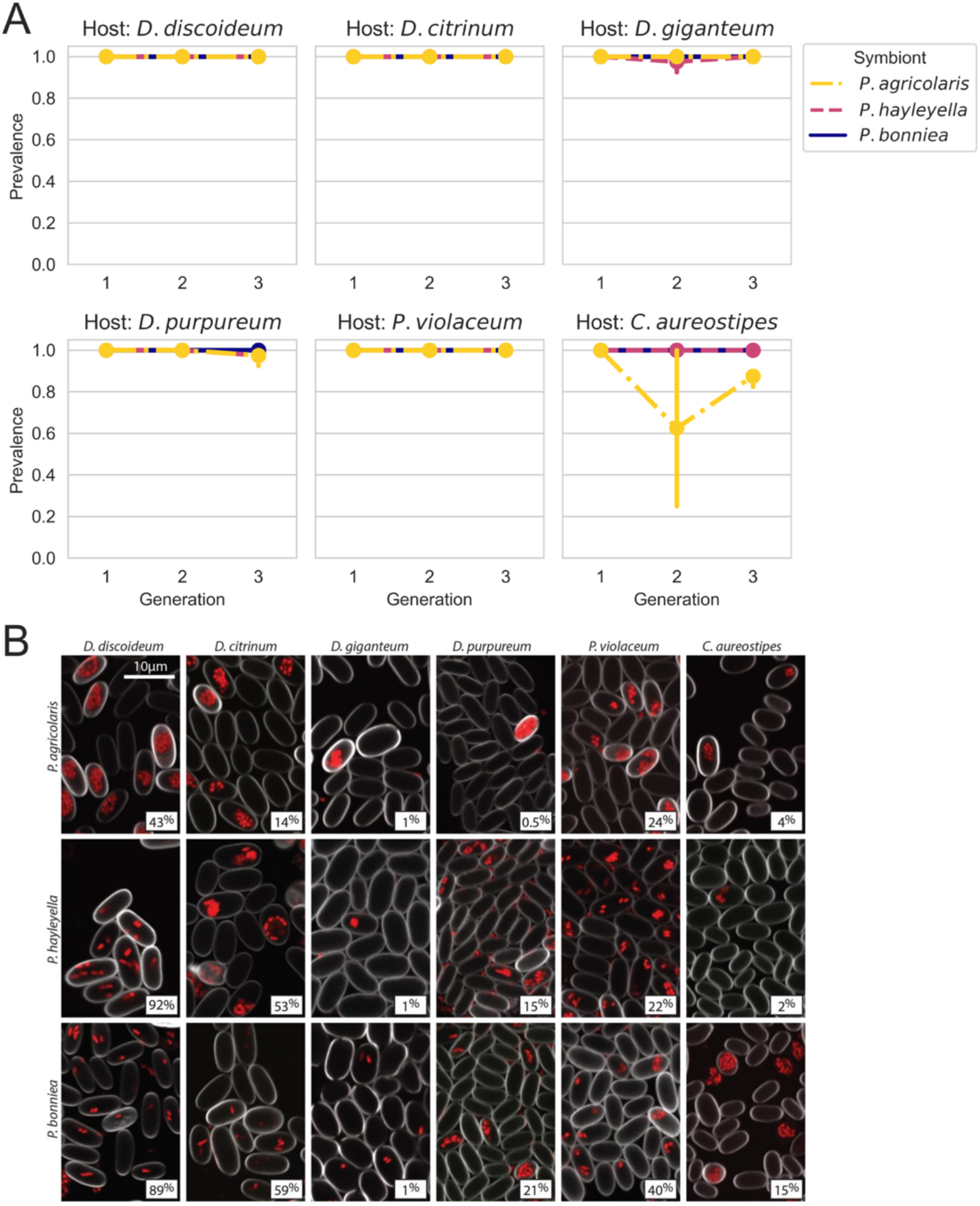
*Paraburkholderia* symbionts form stable and intracellular interactions with a variety of social amoeba hosts upon experimentally mediated dispersal to their local environment. A) The prevalence of *Paraburkholderia* symbionts in each amoeba host species. B) Confocal microscopy images of developed sori contents from indicated amoeba species following exposure to RFP labeled *Paraburkholderia* symbionts. Images show that each *Paraburkholderia* can intracellularly infect (to variable degrees) each host species.

### Intracellular infection frequency and fitness consequences of Paraburkholderia symbionts do not vary with their host’s phylogeny

Our previous experiment suggests that given sufficient dispersal, each *Paraburkholderia* symbiont could stably infect each amoeba species, so we were then interested in the potential role of pre-adaptation in facilitating this dynamic. If the interactions between a symbiont and a host facilitated significant pre-adaptation, one would predict that the fitness consequences of the interaction (for both host and symbiont) in other hosts should be comparable to that of the focal host. Consistent with this prediction, we found that the intra-host fitness of each *Paraburkholderia* symbiont (measured by proportion of host spores infected) did not vary with the phylogenetic distance from their presumed focal host *D. discoideum* (p ≥ 0.2013, see Appendix section 2.1 for full model summaries) (Figure 3A). However, we also found that each symbiont exhibited a reduced ability to infect host spores in a highly diverged host species (*C. aureostipes*), suggesting limitations on pre-adaptation (Figure 3A). Likewise, we found that the host fitness consequences (measured by host spore production) of association with each symbiont did not vary with phylogenetic distance from the presumed focal host *D. discoideum* (p ≥ 0.206, Appendix section 2.2 for full model summaries) (Figure 3B).

**Figure 3.**
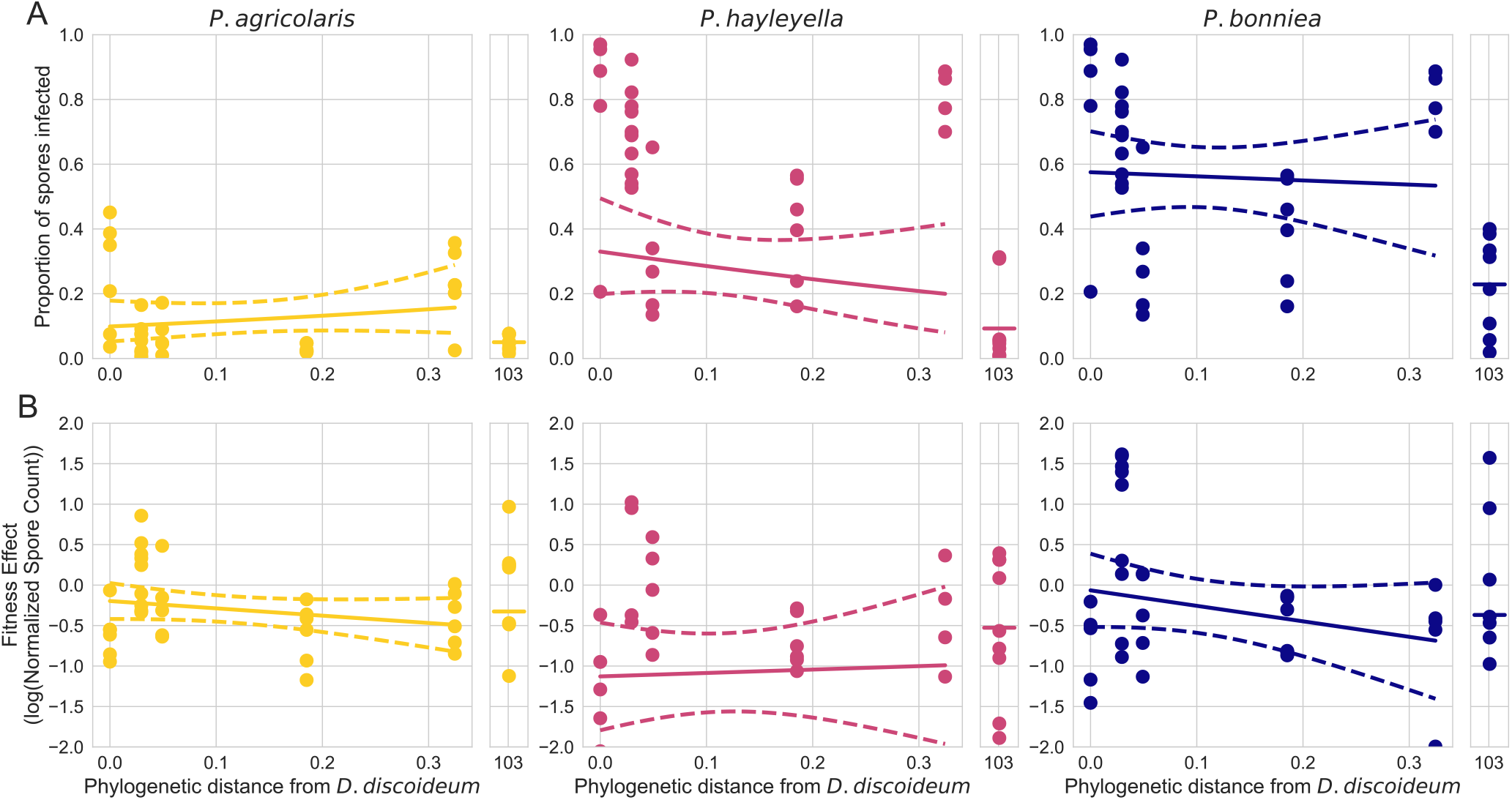
The fitness effects and intracellular infection prevalence of host-symbiont interactions do not vary with increasing phylogenetic distance from their focal host *D. discoideum*. A) The proportion of host spores symbionts were able to infect as a function of phylogenetic distance from *D. discoideum*. B) Log-transformed normalized spore counts as a function of phylogenetic distance from *D. discoideum*. Note that values less than 0 indicate that infected hosts produced fewer spores than an uninfected control. For each panel, sold lines represent the fitted model predictions, dashed lines represent the 95% confidence interval, and points represent the observed data.

### Social amoebae interact with a diverse range of Burkholderiales symbionts in natural populations

During our screenings of natural populations, we found that Dictyostelid social amoebae harbor a diverse range of symbionts within the Burkholderiales order. Specifically, we found 79 unique LepA haplotypes spanning 15 Burkholderiales genera across all host species (Figure 4A). Host species varied in the overall frequency at which they harbored Burkholderiales symbionts. The total prevalence in most host species as between 10% and 20% (Figure 4B). However, Burkholderiales prevalence was over twice as high in *D. discoideum* and *R. minutum* (Figure 4B).

**Figure 4.**
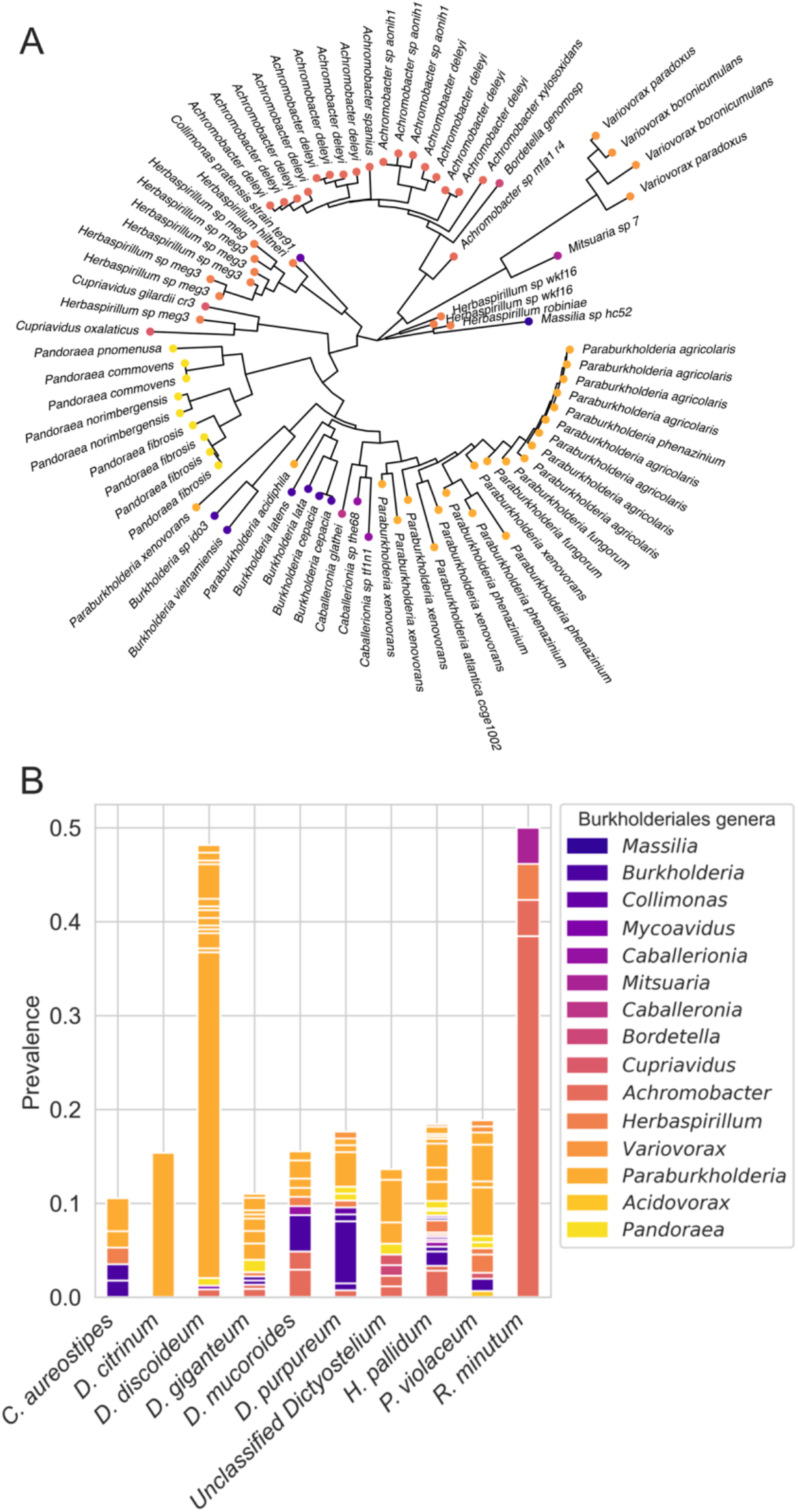
Dictyostelid social amoeba associate with diverse species of Burkholderiales in nature. A) A phylogeny based on the LepA gene of Burkholderiales symbionts found associated with each sampled social amoeba host. Across all sampled amoeba hosts, we identified 79 unique LepA haplotypes from 15 Burkholderiales genera. B) The prevalence of each Burkholderiales haplotype within their respective amoeba host species. Each bar corresponds with a host species, and the y-axis show the prevalence of symbionts within said host species. Note that some bars contain subdivisions, which indicate multiple LepA haplotypes were found within the corresponding genus.

## Discussion

Understanding how ecological processes shape the frequency and distribution of endosymbionts that are segregating in host populations is fundamental for understanding the evolution of endosymbiotic interactions. Here, we found evidence that dispersal limitation plays and important role in shaping the distribution of endosymbioses between social amoeba and their *Paraburkholderia* symbionts. Our survey of natural social amoeba communities showed that patterns of *P. agricolaris* (the only previously described endosymbiont found in our natural survey) diversification were predominately explained by geographic dissimilarity, suggesting the interaction experiences a significant degree of dispersal limitation (Figure 1). We then experimentally mediated symbiont dispersal to host environments to test the importance of dispersal limitation in determining the distribution of these interactions. We found that each previously described *Paraburkholderia* symbiont was able to stably and intracellularly infect each host species at a high prevalence (Figure 2). Furthermore, host-selective dynamics (fitness consequences of the interaction) did not vary with increasing phylogenetic distance from the focal host *D. discoideum* (Figure 3). Finally, we also found that Dictyostelid social amoebae tend to associate with a wide diversity of Burkholderiales bacteria (Figure 4).

Overall, our findings suggest that the endosymbiotic interactions between Dictyostelid amoeba and *Paraburkholderia* symbionts are shaped by a high degree of dispersal limitation, where pre-adaptation facilitates their interactions upon occupation of a shared environment. Therefore, it is possible that limitations on symbiont or host dispersal, rather than specific symbiont functions, explains the variable frequency and often absence of these interactions in natural populations (Haselkorn et al. 2019; DuBose et al. 2022). Laboratory studies have described several possible functions of *Paraburkholderia* symbionts for their amoeba hosts, which include toxin resistance (Brock et al. 2016) and facilitating bacterial food carriage (DiSalvo et al. 2015). However, these symbionts generally have negative fitness consequences (which our findings recapitulated) and field studies have not found evidence that these laboratory-described functions are relevant in natural contexts (DiSalvo et al. 2015; Haselkorn et al. 2019; DuBose et al. 2022). Therefore, it seems unlikely that selective benefits conferred by previously suggested symbiont functions have played an important role in the ecological or evolutionary dynamics of these interactions. Rather, our findings suggest a simpler model where Dictyostelid amoebae happen to ingest *Paraburkholderia* symbionts that are in their local environment, and said symbionts are generally pre-adapted to survive ingestion by amoeba and confer some fitness cost. Our finding that a wide diversity of Burkholderiales symbionts seem to associate with Dictyostelid hosts suggests this model may apply generally to other symbionts besides those that have been previously described. However, we suggest this with the caveat that we did not discern whether these other symbionts could actually persist within the host.

Considering dispersal limitation has the potential to significantly expand our understanding of the evolutionary dynamics of these amoebae-bacteria endosymbioses. For example, even if previously described symbiont functions do occur in nature, their relevance may only be sparsely realized by a small portion of the host population. In other words, dispersal limitation could limit the selective spread of potential beneficial associations, which would also be consistent with the highly variable prevalence seen in nature (Haselkorn et al. 2019; DuBose et al. 2022). It is easy to consider a scenario where the frequency at which a symbiont appears in association with a host is significantly influenced by the frequency at which it disperses to said host’s environment (or vice versa). Theory suggests that at low frequencies, the symbiont would be more subject to ecological drift and would have a greater likelihood of being stochastically lost (Vellend 2010). Our study is limited in that we manipulated symbiont dispersal at a fixed density. However, future studies that examine symbiont colonization/loss in amoeba populations as a function of their immigration density could directly test this hypothesis, which would give useful insight into the incredibly variable symbiont prevalence seen in nature.

Our study is not able to conclude whether limitations on host dispersal, symbiont dispersal, or both is responsible for the observed patterns of dispersal limitation of the interaction. There is evidence that natural Dictyostelid populations are dispersal limited, as genetic differentiation between *D. discoideum* populations has been observed at narrow and broad geographic scales (Douglas et al. 2011; Kuzdzal-Fick et al. 2023). However, less is known of population structure in other Dictyostelid species. Symbionts could also experience limitations on dispersing to host environments. For example, abiotic conditions impact the growth and distribution of Burkholderiales bacteria, including the symbionts studied here (Stopnisek et al. 2014; Brock et al. 2020). Therefore, abiotic conditions could selectively limit the distribution of symbionts, which could in turn limit their dispersal to host environments. It is likely that both factors are at play, but further work would be needed to establish their relative contributions.

Consistent with previous studies, we also found that *Paraburkholderia* symbionts appear pre-adapted to survive and proliferate in a variety of Dictyostelid hosts (Mather et al. 2023). Furthermore, we found that the fitness effects of these interactions did not decline with increasing phylogenetic distance from the focal host *D. discoideum*. This is generally consistent with a previous study that found a variety of *P. bonniea* symbionts strains did not have different consequences on fitness when paired with native and novel host genotypes (Noh et al. 2024). Given the growing body of evidence suggesting a lack of specificity in these interactions, it is possible that *D. discoideum* is not necessarily the focal host as previously suggested. The genomes of *P. bonniea* and *P. hayleyella* symbionts show significant signatures of erosion due to long-term occupation of intracellular host environments (Brock et al. 2020). However, natural surveys have consistently found that they do not frequently associate with *D. discoideum* (Haselkorn et al. 2019; DuBose et al. 2022), and our findings now suggest that they do not frequently associate with other common Dictyostelid species either. Therefore, it is possible that their primary host is not *D. discoideum* or even another closely related species. Rather, it is possible that previous recordings of these associations in nature are simply the product of infrequent dispersal from an alternative focal host’s population into *D. discoideum* populations. Conversely, the *P. agricolaris* genome and physiology is more similar to other free-living *Paraburkholderia* species, and also forms the most frequent associations with Dictyostelid hosts. Therefore, it is possible that its general patterns of association with Dictyostelid hosts are the product of slightly alleviated dispersal limitation relative to the other two symbionts, which could be mediated by its greater competitiveness when free-living (Brock et al. 2020).

More generally, the role that dispersal plays in shaping the frequency of host-symbiont interactions is fundamental for understanding their evolutionary dynamics. In many different systems, there are often discrepancies between symbiont functions described in laboratory studies and the apparent relevance of said functions in nature (Hrček et al. 2016; DuBose et al. 2022). There is a growing appreciation for the role of selection in explaining these complexities, where the net benefit or detriment of a symbiotic interaction changes depending on what stressors are at play in the environment at a given time (Oliver et al. 2014). Our findings suggest it is also important to understand the role of dispersal in limiting or facilitating the spread of potential functions. For example, if a symbiotic interaction becomes beneficial in only a portion of a population that is dispersal limited, it would be more difficult for this benefit to be generalized to the whole population. This issue compounds when considering a given interaction may only be beneficial between hosts and symbionts of particular genetic backgrounds or in particular environments. Therefore, future studies on the interplay between dispersal and other ecological processes have the potential to significantly improve our understanding of the evolution of endosymbioses.

## Acknowledgements

We would like to UCA Biology Department and College of Natural Sciences and Mathematics for funding. Other undergraduates contributing to this work include Erin Golden, Britteny Beruman, Lorrin Hooten, Anthony Barkdull, and Hunter Olsen. We also thank Sydni Rubio for contributing to data collection. This project has been supported by the Arkansas INBRE program, with a grant from the National Institute of General Medical Sciences (NIGMS), P20 GM103429 from the National Institute of Health.

## Data and Code availability

All sequences generated for this manuscript will be available on GenBank upon publication. All code and data associated with this manuscript can be found at https://github.com/gabe-dubose/paraburk_dispersal.

