## Appendix for "The roles of dispersal limitation and pre-adaptation in shaping *Paraburkholderia* endosymbiont frequencies in social amoeba communities"

### 1 Extended Methods

#### 1.1 Culture media and buffers

##### 1.1.1 Hay agar

Ingredients (per liter):

- 15g hay
- 1.5L deionized $H_2O$
- 1.5g $KH_2PO_4$
- 0.62 g $Na_2HPO_4$
- 15g agar

Preparation:
Boil water in solution until infused. Filter hay out of solution and add each ingredient. Autoclave and pour before solution solidifies.

##### 1.1.2 SM/5 agar

Ingredients (per liter):

- 1L deionized $H_2O$
- 2g glucose
- 2g BactoPeptone
- 2g yeast extract
- 0.2g $MgCl_2$
- 1g $K_2HPO_4$
- 1.9g $KH_2PO_4$
- 15g agar

Preparation:
Mix each ingredient, autoclave, and pour before solution solidifies.

##### 1.1.3 KK2 spore buffer

Ingredients (per liter):

- 1L deionized $H_2O$
- 0.67g $K_2HPO_4$
- 2.25g $KH_2PO_4$

Preparation:
Mix ingredients, autoclave.

#### 1.2 PCR primers and protocol

##### 1.2.1 Dictyostelid

D307F: 5'-GTTTGGCCTACCATGGTTGTAA-3'
D862R: 5'-GGGSGTTCATATTGGGGCG-3'

##### 1.2.2 Burkholderiales

Burk3F: 5'-CTGCGAAAGCCGGAT-3'
Burk3R: 5'-TGCCATACTCTAGCYYGC-3'

##### 1.2.3 LepA

LepAF: 5'-CTSATCATCGAYTCSTGGTTCG-3'
LepAR: 5'-CGRTATTCCTTGAACTCGTARTCC-3'

##### 1.2.4 Protocol

95 C (30sec), 55 C (30s), 72 C (45s) x 35 cycles

### 2 Supporting results

#### 2.1 Model summaries for spore infection prevalence as a function of phylogenetic distance from *D. discoideum*

##### 2.1.1 *P. agricolaris*

Table S1: Quasi-binomial model summary

| $C$oefficients | Estimate | Std. Error | t | $Pr(>\|t\|)$ | 95% CI |
|---|---|---|---|---|---|
| Intercept | 0.099 | 0.3497 | -6.320 | 5.72e-07 | 0.05 - 0.17 |
| Distance from *D. discoideum* | 0.83671519 | 1.8480 | 0.884 | 0.384 | 0.11 - 0.99 |

##### 2.1.2 *P. hayleyella*

Table S2: Quasi-binomial model summary

| $C$oefficients | Estimate | Std. Error | t | $Pr(>\|t\|)$ | 95% CI |
|---|---|---|---|---|---|
| Intercept | 0.33 | 0.3501 | -2.021 | 0.0526 | 0.20 - 0.49 |
| Distance from *D. discoideum* | 0.11 | 2.2986 | -0.912 | 0.3695 | 0.001 - 0.91 |

##### 2.1.3 *P. bonniea*

Table S3: Quasi-binomial model summary

| $C$oefficients | Estimate | Std. Error | t | $Pr(>\|t\|)$ | 95% CI |
|---|---|---|---|---|---|
| Intercept | 0.56 | 0.2815 | 1.078 | 0.290 | 0.44 - 0.70 |
| Distance from *D. discoideum* | 0.37 | 1.9088 | -0.271 | 0.788 | 0.014 - 0.96 |

#### 2.2 Model summaries for host fitness response to infection as function of phylogenetic distance from *D. discoideum*

##### 2.2.1 *P. agricolaris*

Table S4: Quasi-binomial model summary

| $C$oefficients | Estimate | Std. Error | t | $Pr(>\|t\|)$ | 95% CI |
|---|---|---|---|---|---|
| Intercept | -0.1968 | 0.1133 | -1.737 | 0.0926 | -0.43 - 0.035 |
| Distance from *D. discoideum* | -0.9041 | 0.6920 | -1.306 | 0.2013 | -2.32 - 0.51 |

##### 2.2.2 *P. hayleyella*

Table S5: Quasi-binomial model summary

| $C$oefficients | Estimate | Std. Error | t | $Pr(>\|t\|)$ | 95% CI |
|---|---|---|---|---|---|
| Intercept | -1.1282 | 0.3396 | -3.322 | 0.00243 | -1.82 - -0.43 |
| Distance from *D. discoideum* | 0.4243 | 2.0425 | 0.208 | 0.83690 | -3.753070 - 4.6015832 |

##### 2.2.3 *P. bonniea*

Table S6: Quasi-binomial model summary

| $C$oefficients | Estimate | Std. Error | t | $Pr(>\|t\|)$ | 95% CI |
|---|---|---|---|---|---|
| Intercept | -0.06471 | 0.23094 | -0.280 | 0.781 | -0.54 - 0.41 |
| Distance from *D. discoideum* | -1.91432 | 1.48 | -1.3 | 0.206 | -4.94 - 1.12 |
